# Male social dominance affects access to mates but not female mate choice in a livebearing fish

**DOI:** 10.1101/2025.10.16.682567

**Authors:** António Gonçalves, Charel Reuland, Léa Daupagne, David Wheatcroft, Niclas Kolm, John L. Fitzpatrick

## Abstract

Male-male competition and female mate choice are often assumed to act in concert, with male social dominance serving as a reliable cue in female choice reflecting overall male quality. However, few studies have explicitly disentangled the effects of male competition and female mate choice, leaving gaps in our understanding of how these processes interact to shape male fitness. Here, we experimentally separated the effects of male-male competition and female mate choice in the pygmy halfbeak *Dermogenys colletei*, a small freshwater fish, to test whether i) male social dominance drives access to females in the presence and absence of direct male-male competition, and ii) females prefer socially dominant males. Our results show that dominant males monopolized access to females, spending more time courting females and engaging in more courtship behaviors. Additionally, female halfbeaks did not show a preference for dominant males, providing no evidence that females use male dominance when assessing males. Overall, these findings highlight that when male-male competition facilitates mate monopolization there may be less scope for pre-copulatory female mate choice.

**Lay summary:** Social dominance helps male pygmy halfbeaks gain access to mates, as dominant males court more and spend more time with females. However, females do not prefer dominant males, suggesting that competition between males, rather than female choice, may determine mating success in this species. This highlights how social dominance can shape mating opportunities even when female preferences are weak.

## 1 Introduction

An individual’s reproductive success is shaped by both competition among members of the same sex for mating opportunities and mate choice by members of the opposite sex (Darwin, 1871; Kokko et al., 2012). In species with Darwinian sex roles, males typically exhibit greater variance in reproductive success than females (Janicke et al., 2016), and consequently often experience intense male-male competition for access to mates. In turn, the sex limited by their lower potential reproductive rate, typically females due to their higher investment into offspring, commonly benefits from selecting high-quality mates (female mate choice). Intra-sexual competition and mate choice are expected to generate directional selection for traits that are correlated with success in either process, leading to the evolution of exaggerated traits, such as weapons and larger body sizes used to resolve competitive interactions (Emlen, 2008; Hardy and Briffa, 2013; McCullough et al., 2016). However, there is ample scope for interactions between these mechanisms of sexual selection, with male-male competition and female mate choice potentially acting in concert, in opposing directions, or independently (Berglund et al., 1996; Wiley and Poston, 1996; Wong and Candolin, 2005). Yet, despite its potential to shape the strength and direction of sexual selection (Hunt et al., 2009), interactions between male-male competition and female mate choice remain poorly understood (Wong and Candolin, 2005; McCullough et al., 2016).

Male-male competition and female mate choice have traditionally been viewed as mutually reinforcing mechanisms of sexual selection (Berglund et al., 1996; Wiley and Poston, 1996; Wong and Candolin, 2005). Traits that provide males with a competitive advantage in male-male competition may also represent a reliable indicator of male quality (Berglund et al., 1996; Khazraıe and Campan, 1999). Females who select dominant males as mating partners may therefore gain either direct fitness benefits, such as access to better resources or protection, or indirect benefits, such as superior genetic traits for their offspring (Berglund et al., 1996; Wong and Candolin, 2005; Hunt et al., 2009). Female preference for dominant males has been recorded for a variety of species (reviewed in Wong and Candolin, 2005). Such female preference can be shaped by females gaining information about social dominance by observing interactions between males (Doutrelant and McGregor, 2000). Competitive interactions between males are a conspicuous form of social information, akin to other information available in the social environment that influences mate choice decisions (e.g., other individuals mating, Dugatkin, 1996, presence of rivals, Plath and Bierbach, 2011). The reproductive outcomes associated with female preference typically translate into higher fitness for dominant males (Bulger, 1993; Rus Hoelzel et al., 1999; Baxter et al., 2015; Montana et al., 2022). However, distinguishing whether reproductive benefits observed in dominant males arise from active female preference or male-imposed constraints remains challenging without experimental manipulation. Indeed, in many species, males that are more successful during male-male competition can monopolize access to females, thereby constraining female choice by preventing interactions between females and subordinate males and reducing the overall pool of potential mates (Wong and Candolin, 2005).

Experimental studies examining the interaction between male-male competition and female mate choice have produced mixed results. In some cases, females prefer to associate with socially dominant males after assessing visual and olfactory cues associated with agonistic interactions between males, as shown in fighting fish (*Betta splendens*, Doutrelant and McGregor, 2000), red swamp crayfish (*Procambarus clarkii*, Aquiloni et al., 2008; Aquiloni and Gherardi, 2010), and black-capped chickadees (*Parus atricapillus*, Mennill et al., 2002). In contrast, other species show no effect of male-male competition on female preference, including the Pacific blue-eye fish (*Pseudomugil signifier*, Wong, 2004) and the jumping spider (*Thiania bhamoensis*, Chan et al., 2008). Some species even exhibit reversed patterns, where females prefer subordinate males after observing male social interactions, as observed in the Mexican livebearing fish (*Poecilia mexicana*, Bierbach et al., 2013) and Japanese quail (*Coturnix japonica*, Ophir and Galef Jr, 2003). These findings suggest that the influence of male-male competition on female mate choice is complex, context-dependent and species-specific, highlighting the need for further research to clarify how these mechanisms interact.

In this study, we investigate how male-male competition influences access to mates and female mate choice in the pygmy halfbeak *Dermogenys collettei*, a small, internally fertilizing, freshwater fish native to south-east Asia (Meisner, 2001; Greven et al., 2010). Halfbeaks live in loosely organized mixed-sex groups, where intra- and inter-sexual interactions are frequent (Greven et al., 2010; Devigili et al., 2021). Males compete with rival males for access to females, often directing behavioural threat displays to rivals and using their elongated lower jaws (called beaks) as an armament during competitive interactions. Furthermore, males are vigorous courters (Devigili et al., 2021), and females can discriminate and choose males based on the amount of red fin coloration displayed (Reuland et al., 2020; McNeil et al., 2021). In highly competitive scenarios (i.e. when agonistic interactions between males are frequent), dominant males exhibit superior ejaculate quality compared to subordinate males (Reuland et al., 2021), hinting at the potential benefits of female preference for dominant males. Here, we experimentally examine i) how male social dominance influences access to females in the presence or absence of direct physical interactions between males and ii) how the outcome of male-male competition influences female choice. We predicted that i) dominant males would have greater access to mates, but only when they could physically interact with subordinate males, and ii) female preference would be based on male social dominance, with females preferring dominant over subordinate males.

## 2 Methods

### 2.1 Study populations and rearing conditions

Halfbeaks used in this study were sexually mature, wild-caught (Experiment 1) or lab-reared descendants of a wild-caught population (Experiment 2) from the Johor State of Malaysia (Johor River for Experiment 1 and Tebrau River for Experiment 2). The different sources of fish were a consequence of the availability of fish from a commercial exporter. While we have no *a priori* reason to expect behavioural differences between these sources, population-level differences cannot be completely excluded. To avoid confounding effects of source within experiments, all analyses were restricted to fish derived from a single source. Fish were reared in the laboratory using a standard set of husbandry practices. Fish were maintained on a 12:12 light:dark cycle at 26-27°C and fed daily a mixture of either ground flake food, freshly hatched Artemia, *Drosophila melanogaster*, or mosquito larvae (*Chaoborus crystallinus*). Aquaria contained ∼2 cm of gravel, were oxygenated, and contained live or artificial plants. Sexually mature fish were kept in mixed sex 160L aquaria in groups of 10-25 individuals. To generate lab-reared descendants of wild-caught fish, gravid females were placed in 7.5L tanks and monitored daily until they produced offspring (i.e., fry). Alternatively, fry were removed from stock tanks whenever they were observed. Fry were then separated from adults to prevent cannibalism and kept in 5-7.5L tanks with up to five other fry. The onset of sexual maturity was determined by observing a thickening of the male’s andropodium, the modified set of five fin rays that males use to transfer sperm to female (Meisner and Burns, 1997; Greven et al., 2010). At the onset of sexual maturity (∼4 months), females and males were separated into either mixed sex or sex-specific tanks (to maintain their mating status as ‘unmated’) and reared in 160L aquaria. Three days before each experiment, males and females were removed from their stock tanks, photographed for phenotypic trait quantification (see 2.2.1), and housed individually in 7.5 L tanks labeled with individual name tags to allow identification throughout the experiment.

### 2.2 Experimental overview

We performed two experiments to assess how male social dominance influences reproduction in halfbeaks. In Experiment 1, we tested the hypothesis that male-male social dominance influences access to mates by examining the link between male social dominance and courtship behaviours directed towards females in the presence or absence of direct physical interaction between males. In Experiment 2, we tested the hypothesis that male-male social dominance influences female mate preferences.

#### 2.2.1 Experiment 1: Does male social status influence access to mates?

To assess the effect of male social dominance on access to mates, we placed two unfamiliar males and two randomly selected females in an experimental tank (42 x 24.5 x 17.5 cm), which was divided differently depending on the experimental treatment described below. Males were closely size-matched within dyads, with mean absolute differences in body length of 1.32 ± 1.18 mm across pairs. All tanks were filled to a depth of ∼12 cm and contained a ∼1cm layer of gravel. All experimental treatments began in the same way. First, each experimental tank was partitioned into two compartments using an opaque barrier. Both males were placed on one side of the barrier (24.5 x 24.5 x 17.5 cm), while both females were placed on the other side (30 x 24.5 x 17.5 cm). Within the male compartment, an additional opaque divider was used to further separate the males, confining each male to an individual area (12 x 24.5 x 17.5 cm) with no visual access to one another at the start of the experiment. In contrast, the two females could interact freely with one another but had no visual access to either male. Fish were given a 30-minute habituation period after being placed in their respective areas of the experimental tank. Following habituation, the experiment proceeded in two phases: a male-male competition phase and a mate interaction phase.

##### Male-male competition phase

During the male-male competition part of the experiment, the opaque barrier separating the two males was lifted and the males were allowed to interact for 30 minutes. A total of 74 males were used, forming 37 dyads. Male-male competitive interactions are common in halfbeaks, with a range of agonistic behaviours used during contests with rivals. Contests typically start with both males swimming parallel to each other and *gill flaring*, a behaviour which consists of repeatedly opening and closing their gills usually with their beak open. During these standoffs, males also *head shake* by jerking their bodies in an S pattern. Contests can include biting and chasing (Devigili et al., 2021; Reuland et al., 2021). Occasionally, males interlock their beaks together, a behaviour known as *beak wrestling* (Greven et al., 2010). Following a rival’s aggressive display, an individual may be displaced, turning quickly and swimming away, increasing the distance between himself and its aggressor (Reuland et al., 2021). Male dominance within each dyad was scored during this interaction period by recording displacement behaviours, where one male (the aggressor) approaches the other or performs an agonistic behaviour (e.g., gill flares, head shakes, bites, chasing), making the targeted male distance himself from the aggressor male. These displacement behaviours were used to create a dominance index, calculated as:

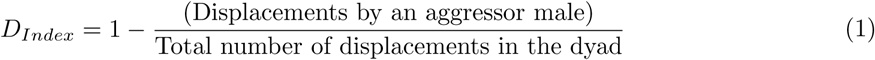

Dominance index values range from 0 to 1, with 0 meaning a male was completely subordinate (i.e., males were only displaced by rival males) and 1 completely dominant (i.e., males only displaced rival males, Reuland et al., 2021). Following Reuland et al. (2021), males were classified as socially dominant in their dyad if they had a dominance score *>* 0.7 (i.e., males displaced their rival in *>* 70% of interactions). Dominant males had an average ± SE dominance index score of 0.96 ± 0.04, while the dominance index score of subordinate males was 0.05 ± 0.04. Importantly, the opaque barrier partitioning the males from females was left in place throughout the male-male competition part of the experiment to prevent females from directly observing male-male interactions.

##### Mate interaction phase

Following the male-male competition phase of the experiment, the opaque barrier separating males and females was lifted. In n=13 trials, both the dominant and subdominant males (26 males in total) were allowed to freely interact with both females and one another for 30 minutes (henceforth called *mate interactions with male-male competition* trials). In n=24 trials, only one of the two males was allowed to interact with the two females, while the other male was placed behind a transparent partition (henceforth called *mate interactions without male-male competition* trials). The partition prevented the focal males from physically interacting with the males confined by the transparent partition (i.e. the stimuli males) during the mate interaction part of the experiment. However, the transparent partition allowed the passage of visual signal, and by drilling several holes into the partition, we allowed the exchange of chemical cues between males and females. Thus, in the *mate interaction* trials either *with* or *without male-male competition*, the number of fish visually present remained constant, ensuring a consistent adult sex ratio across trials, while the potential for male-male competitive interactions to affect access to mates varied. In the *mate interactions without male-male competition* trials, males were randomly assigned to either the focal or stimuli male role (i.e., males were assigned blind to their social dominance scores from the previous part of the experiment). As a result, the sample size was n=14 dominant males and n=10 subordinate males acting as focal males (and conversely n=10 dominant males and n=14 subordinate males as stimuli males).

In *mate interactions with* and *without male-male competition*, we recorded the total amount of courtship behaviours males directed towards either female. Courtship behaviours include *circling*, where the male swims in a semi-circular pattern around the female, *swimming under*, where the male positions himself under the female with his head located below the female’s genital pore, and *mating*, where the male rapidly (∼40-80 ms) twists his body in an attempt to use his andropodium to transfer sperm to the females’ genital pore (Greven et al., 2010). Males frequently attempt to swim under females, with some attempts being successful and others being considered unsuccessful when females swim away as males attempt to swim under. We treated the total number of *swimming under attempts* as a measure of male sexual interest. Other courtship behaviours present in the male’s repertoire (e.g., *nipping*, where males rapidly open and close their beaks while swimming under the female, and *checking* where males use their beak to make physical contact with the female) were difficult to observe in our experimental setup and were therefore not considered. We recorded the total number of courtship behaviours (circling, swimming under attempts and mating attempts, henceforth called total courtship count) and the total amount of time males spent swimming under each female.

All assays were recorded by the use of GoPro 10 and videos were analysed with the use of the software BORIS (version 8.24.1; Friard and Gamba, 2016).

##### Quantifying phenotypic traits

As both male and female phenotypes could influence mate choice in halfbeaks (Reuland et al., 2020; Ogden et al., 2020; Reuland et al., 2021; McNeil et al., 2021), we quantified relevant phenotypic traits of all fish used in the experiment. We placed the fish inside an aquarium (27 x 17.5 x 15.5 cm), filled with conditioned water, with the aquarium itself positioned inside a photography chamber under standardized light conditions and fitted with a scale. Fish were positioned against the side of the aquarium with help from a plastic sheet with a groove deep enough to hold them in place for the pictures and not cause them any physical harm. The fish were only held in this position for a few seconds while the pictures were taken. The photos were taken with a Canon EOS 800D camera and were used to measure body length (mm), measured from the anterior side of the eye to the caudal peduncle (Fernlund Isaksson et al., 2022), beak length (mm) and total area of red coloration on the anal, dorsal and caudal fin (mm^2^) using the polygon selection tool in ImageJ (version 1.54; Schneider et al., 2012). For stimuli females, the females’ ventral side was photographed by placing females (for *<* 1minute) in a clear plastic photography container (75 x 50 x 25 mm) filled with water from their own tank. The container was placed on top of a plexiglass sheet held in place above the same Canon camera that was used to take pictures from the side of the fish. From these pictures, female body length (mm) and gravid spot area (mm^2^) was measured using ImageJ.

#### 2.2.2 Experiment 2: Does male social status influence female mate preferences and female mating behaviour?

To assess if male social dominance influences female mate preferences, we performed mate choice assays in dichotomous choice tanks (45 x 25 x 20 cm), consisting of one main chamber (45 x 15 x 20 cm) and three smaller stimulus chambers (each 15 x 10 x 20 cm). All chambers were separated by removable transparent and opaque dividers, allowing for changes in chamber configurations throughout the experiment. A focal virgin female was placed in the main chamber. While virgin females are commonly assumed to be less choosy than mated females (Jennions and Petrie, 2000; Kokko and Mappes, 2005), a recent meta-analysis found no evidence that mating status influences choosiness in females (Richardson and Zuk, 2023). However, because experience with members of the opposite sex and exposure to courtship behaviours can influence female mating behaviours (Coleman et al., 2004; Dukas, 2005; Bailey and Zuk, 2008), we used virgin females that had visual access to 70 L tanks housing 10 males and 10 females for 11-12 days before entering the experiment. Two unfamiliar males were placed randomly in either the right or left stimuli chambers of the dichotomous choice tanks. Males were photographed three days before entering the experimental tank (as described in section 2.2.1). All fish were given one hour to habituate in the experimental chambers, during which time opaque barriers were in place to prevent the fish from visually interacting with one another.

Following the habituation period, the opaque barriers were lifted and females were allowed to freely explore the experimental chamber and visually interact with both stimulus males for one hour. During this period, the stimulus males remained separated from one another by opaque dividers. This phase was used to quantify *baseline female preference*, allowing females to assess both males in the absence of male–male interactions.

After this baseline preference phase, the opaque barriers separating the stimulus males was removed, allowing the males to freely interact with each other for one hour while the female remained in the main chamber and could observe their interactions. This phase provided females with the opportunity to observe male–male competition, hereafter referred to as the *eavesdropping phase*. During male-male interactions, we recorded the number of times a male displaced the other male and calculated the dominance index score for each male (as above). We also recorded the number of agonistic behaviours displayed (see *mate competition phase* section above) to assess the overall aggression of males.

Following the interaction period, males were guided back to the stimuli chambers and the opaque barriers between the stimuli chambers were re-erected. We then assessed female preference for one hour, during which time they could move freely in the main chamber and visually interact with both males. This phase measured female preference *after observing male–male interactions*.

During both the baseline preference phase and the post-interaction preference phase, the duration of time a female spent within 5 cm in front of each stimulus male’s chamber (termed the *association zone*; total dimension: 15 x 5 cm), was recorded. The time females spend in such association zones is commonly used as a measure of mating preference that is predictive of female preference during mating interactions in many species (Bischoff et al., 1985; White et al., 2003; Walling et al., 2010) and was previously used for halfbeaks (Reuland et al., 2020; McNeil et al., 2021).

In total, 64 dichotomous mate choice trial were conducted. Five trials were excluded from the final analysis because females failed to visit both association zones during the one-hour exploration phase, preventing assessment of baseline interest in both males. Additionally, 13 dyads were omitted as, in each dyad, one male did not displace their rival in *>* 70% of interactions. leading to an unclear dominance hierarchy (scores between 0.52 and 0.69). This reduced the final sample size for analysis to n = 46. All tanks were recorded from above using a Point Grey Grasshopper 3 4.1 megapixel camera (Tele-dyne FLIR LLC, Wilsonville, United States) with Fujinon CF25HA-1 lens (Fujifilm, Minato, Japan). Recorded videos were scored using BORIS (version 8.24.1; Friard and Gamba, 2016).

### 2.3 Statistical analysis

Statistical analyses were carried out using R version 4.3.1 (R Core Team, 2023). Linear and generalized linear mixed effects models (LMMs and GLMMs) were performed using the *lme4* (Bates et al., 2015) and *glmmTMB* (Brooks et al., 2017) packages.

#### 2.3.1 Experiment 1: Does male social status influence access to mates?

To examine whether male access to females was influenced by male dominance status, female body size or gravid spot size, LMMs and GLMMs were used. Two metrics reflecting male mating behaviours (duration of swimming under and total courtship count) were included as response variables. The duration of swimming under the female was log-transformed to obtain a normal distribution and assessed using LMMs. Raw data inspection showed a right-skewed distribution with a substantial proportion of zero observations (approximately 10% zeros across trials) for courtship behaviours; accordingly, these were analysed using GLMMs fitted with a negative binomial distribution for count data. Model diagnostics (Pearson dispersion ratio) indicated adequate model fit, with no evidence of overdispersion. Male identity was incorporated as a random effect in each model to account for repeated measures, as each male contributed two observations per trial (one per female). Interactions between male dominance status and trial type (i.e. mate interactions with or without male-male competition) were also fitted to additionally test whether dominance effects differed between experimental settings. These full models constituted the primary framework for hypothesis testing (Table 1). Additional models fitted separately within each trial type are presented in the Supplementary Material (Table A1) to illustrate context-specific patterns. Other two-way interactions were initially included in all models, but were excluded from the final models as they were non-significant.

**Table 1:** Effect of male social dominance and female traits on male access to females.

| Response variable | N | Predictors | Estimates<br>[95% CI] | SE | Test<br>Statistic | P-values |
| --- | --- | --- | --- | --- | --- | --- |
| (a) *Swimming under (s)* | 100 | Dominance (Dominant) | 1.40 [0.13;2.67] | 0.67 | 2.10 | **0.04(\*)** |
|  |  | Female body size | 0.53 [-0.52;1.58] | 0.55 | 0.96 | 0.34 |
|  |  | Gravid spot size | 0.07 [-0.11;0.25] | 0.09 | 0.73 | 0.47 |
|  |  | Trial type (No competition) | 1.04 [-0.32;2.39] | 0.71 | 1.45 | 0.15 |
|  |  | Dominance x trial type | -0.85 [-2.69;0.99] | 0.97 | -0.88 | 0.38 |
| (b) *Total courtship counts* | 100 | Dominance (Dominant) | 1.16 [0.17;2.18] | 0.50 | 2.31 | **0.02(\*)** |
|  |  | Female body size | -0.20 [0.95;0.55] | 0.38 | -0.54 | 0.59 |
|  |  | Gravid spot size | 0.05 [-0.06;0.16] | 0.06 | 0.96 | 0.34 |
|  |  | Trial type (No competition) | 1.04 [0.02;2.12] | 0.53 | 1.96 | 0.05 |
|  |  | Dominance x trial type | -0.50 [1.94;0.93] | 0.72 | -0.69 | 0.49 |

We tested the effect of male social dominance and female traits on the time each male spent swimming under each female (a) and on all discrete courtship behaviors (b) using linear mixed effects models (LMMs) and generalized linear mixed effects models (GLMMs), in 13 *mate interactions with male-male competition* trials and in 12 *mate interactions without male-male competition* trials. The interaction between dominance and trial type (with vs without male-male competition) was additionally tested. The total number of pairs assessed (N) is presented, along with the standard error (SE), test statistic (z) and *P-value* for each effect. Significant *P-values* are shown in bold.

#### 2.3.2 Experiment 2: Does male social status influence female mate preferences and mating behaviour?

To assess whether female preference was influenced by male social dominance, the strength of preference (SOP) was calculated relative to the dominant male in each dyad as follows:

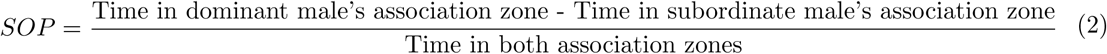

SOP values range from -1 to 1. Positive values indicate a preference for the dominant male, whereas negative values indicate a preference for the subordinate male. A value of 0 indicates equal association time with both males. SOP values were calculated separately for the baseline preference phase (before males interacted, *SOP_baseline_*) and the post-interaction preference phase (after females had observed male–male competition, *SOP_post_*).

To determine whether observing male–male interactions influenced female preferences, we calculated the change in preference for the dominant male between the baseline and post-interaction phases:

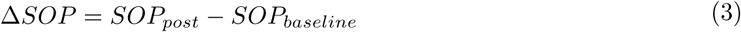

Positive Δ*SOP* values indicate that females increased their association time with the dominant male after observing male–male interactions, whereas negative values indicate a shift toward the subordinate male. A one-sample t-test was used to assess whether the mean Δ*SOP* differed significantly from zero.

Finally, to examine whether morphological differences between stimulus males influenced changes in female preference, SOP was then calculated for the male initially placed in the left stimuli chamber at the start of each trial (i.e., an arbitrary male) as follows:

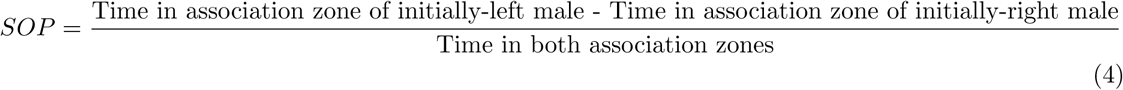

A SOP closer to - 1 indicates a preference towards the right male, while a SOP closer to 1 indicates a preference towards the left male. A SOP of 0 indicated an unbiased mating preference. To assess morphological differences between the left and right stimulus males, we compared their body length, beak length, and total area of red coloration. This was done by subtracting the right male’s measurements from those of the left male. SOP for the left male scores were used as the response variable in a single LMM. The difference between body length, beak length, and red area of the two stimulus males were included as fixed factors, along with male dominance and experimental phase (baseline vs post-interaction). We included tank identity as a random factor to account for potential variation among experimental tanks. Two-way interactions between experimental phase and predictor variables were initially included in the model, but were sequentially excluded from the final models as they were non-significant. Because females could potentially infer male dominance from morphological cues prior to observing male–male interactions, we additionally tested whether dominant males differed from subordinate males in body length, beak length, or red coloration. For each male dyad, we calculated trait differences between the dominant and subordinate male and used one-sample t-tests to determine whether mean differences differed from zero (Table 2).

**Table 2:** Effect of male social dominance and male traits on female mate preference and choice.

| Response variable | N | Predictors | Estimates<br>[95% CI] | SE | *t* | *P-values* |
| --- | --- | --- | --- | --- | --- | --- |
| **Female mate preference** |  |  |  |  |  |  |
| *SOP*<sub>left male</sub> | 46 | Phase (Baseline) | -0.07 [-0.15;0.27] | 0.11 | -0.67 | 0.50 |
|  |  | Dominance (Subordinate) | 0.17 [-0.05;0.41] | 0.12 | 1.44 | 0.15 |
|  |  | Male body length | 0.02 [-0.05;0.09] | 0.03 | 0.51 | 0.61 |
|  |  | Male beak length | 0.01 [-0.16;0.17] | 0.09 | 0.10 | 0.92 |
|  |  | Male red coloration | 0.01 [-0.20;0.22] | 0.11 | 0.08 | 0.94 |

We tested the effect of experimental phase, male social dominance and morphological differences between stimulus males on the strength of preference for the left male scores (an arbitrary male) using a linear mixed effects model (LMM) in dichotomous mate choice trials. The total number of trials (N) is presented, along with the standard error (SE), test statistic (*z* and *P-value* for each effect. Significant *P-values* are shown in bold.

### 2.4 Ethical Note

All procedures in this study were approved by the Stockholm Animal Research Ethical Permit Board (permit number 3967-2020). The potential harm to the animals was limited and primarily related to routine handling and temporary housing in experimental aquaria. Male–male interactions were allowed to occur as part of the experimental design, as such competitive behaviours are common in this species under natural conditions. Interactions were limited in duration and monitored throughout the trials.No injuries or mortality were observed during the experiments. Following the completion of each experiment, all individuals were returned to 160 L social holding tanks. Social groups were carefully observed after reintroduction, and no signs of distress or mortality were detected. Normal levels of social behaviour were typically restored within approximately one hour.

## 3 Results

### 3.1 Experiment 1: Male social dominance influences access to mates

Dominant males swam under females on average 2.1 times longer than subordinate males (Dominants: 97.94±28.01 s, Subordinates: 45.97±15.10 s; mean±SE) (LMM; Table 1, a, Figure 1) and directed 2.0 times more total courtship behaviours towards females (Dominants: 23.08±3.73 behaviours; Subordinates: 11.77±2.23 behaviours) (GLMM; Table 1, b, Figure 1). However, dominance effects did not significantly differ between trial types (i.e. *mate interactions with* versus *without male-male competition* trials, Table 1). Additionally, the duration of time males spent swimming under females and the total courtship counts did not significantly differ based on female body size or the size of the gravid spot (LMM; Table 1, a, b).

**Fig. 1:**
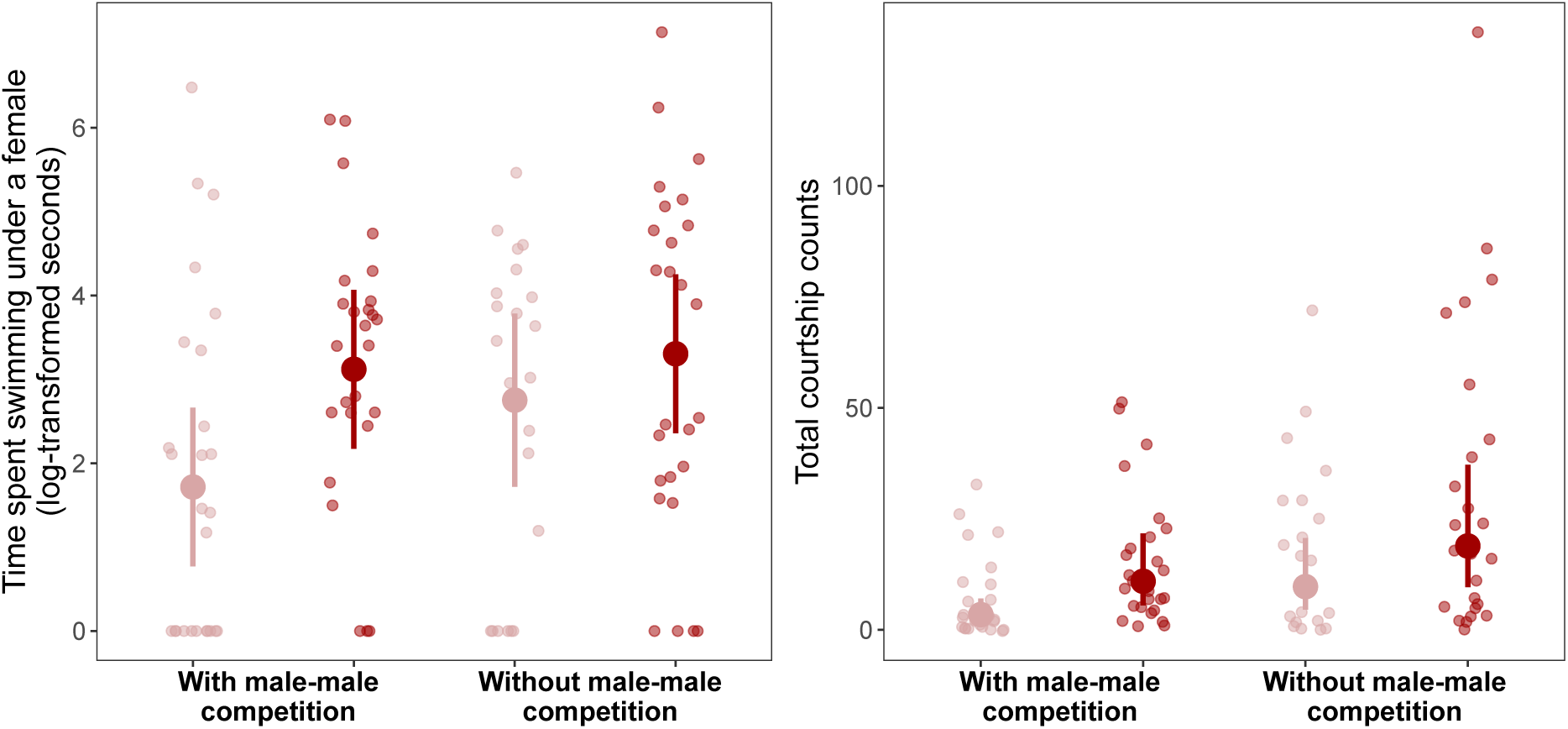
Duration of time (seconds, log-transformed) that each male spent swimming under each female and number of courtship behaviours (i.e. circling, swimming under attempts and mating attempts) performed by each male towards each female during the 30-minute observation period. Subordinate males are shown in light red; dominant males in dark red. Small points represent raw observations, and large dots show model-predicted means ± 95% confidence intervals.

**Fig. 2:**
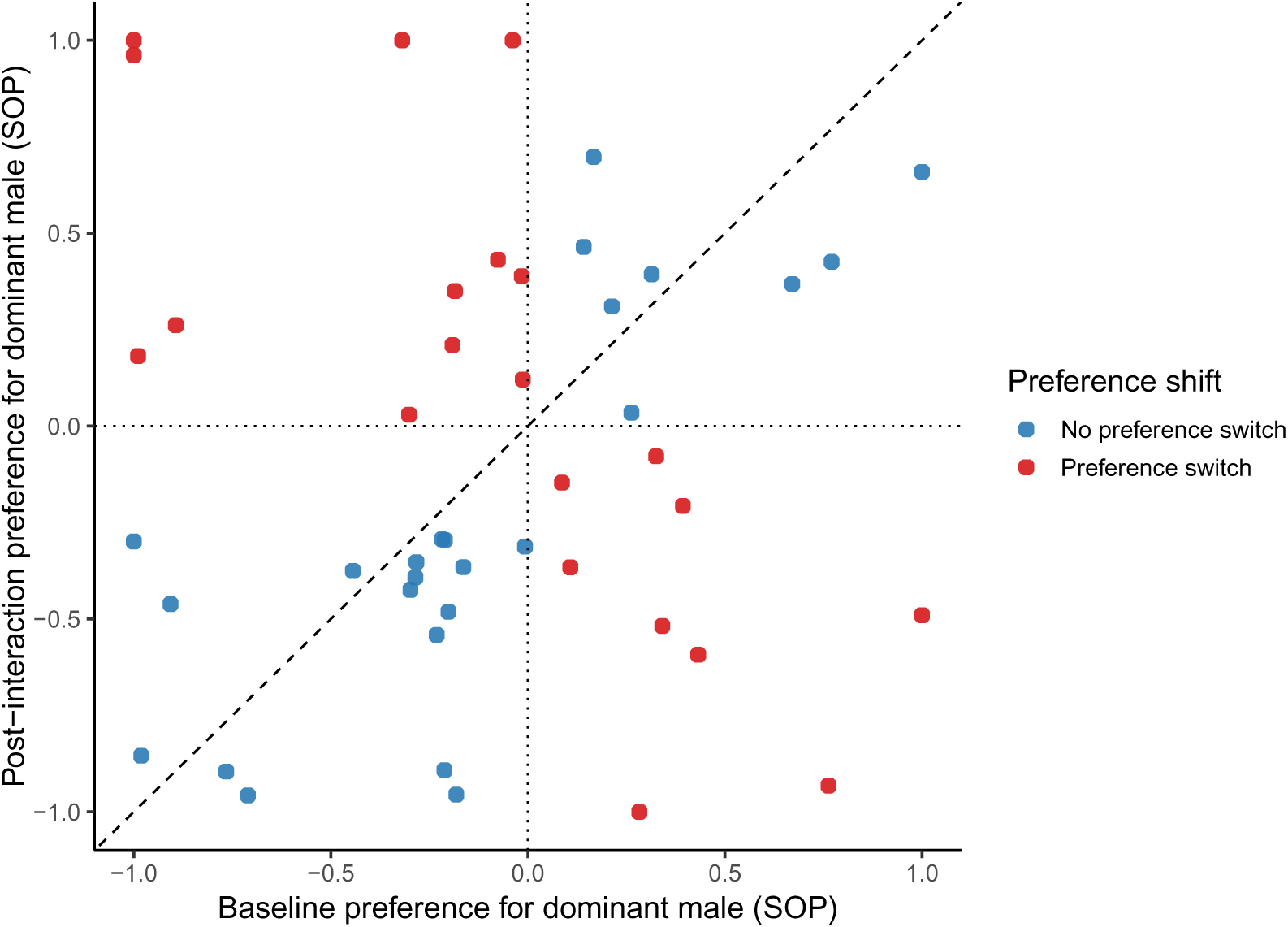
Female preference for dominant males before and after observing male–male competition. Strength of preference (SOP) for the dominant male is shown for the baseline phase (before males interacted) and the post-interaction phase (after females observed male–male interactions). Each point represents one mate choice trial. The dashed line represents no change in preference between phases (Δ*SOP* = 0). Points above the dashed line indicate an increased preference for the dominant male after competition, whereas points below the line indicate a decrease. Red points indicate trials in which females switched preference between males.

### 3.2 Experiment 2: Male social dominance does not influence female mate choice

Females did not alter their preference for dominant males after observing male–male interactions. The change in strength of preference for the dominant male (Δ*SOP*) between the baseline and post-interaction phases did not differ significantly from zero (one-sample t-test: *t*_45_ = 0.13, *p* = 0.90; mean Δ*SOP* = 0.01, 95% CI = [−0.21, 0.24]). This indicates that females did not increase or decrease their association time with dominant males after observing male–male competition.

Consistent with this result, female strength of preference (SOP) was not influenced by experimental phase, male dominance status nor any of the morphological traits (i.e., body length, beak length, red coloration) considered in our analyses (Table 2). Thus, female halfbeaks appear to exhibit an unbiased mating preference with respect to both male social dominance and the morphological traits examined.

## 4 Discussion

In this study, we experimentally examined whether socially dominant males in the pygmy halfbeak gain increased access to females as a result of male-male competition, female mate choice, or a combination of these factors. We first found that dominant males courted females more frequently than socially subordinate males, demonstrating that social dominance confers greater access to potential mates. However, experimental tests of female mate choice indicated that this increased courtship by dominant males was not driven by female preference as females showed no preference for males based on social dominance. Together, these findings suggest that male-male competition plays a stronger role than pre-copulatory female preference in determining male access to females in pygmy halfbeaks.

Social dominance offers a well-known route to securing fitness in a wide range of species (e.g. in fish, Deaton, 2008; Paull et al., 2010; Weir et al., 2004; in birds, Nelson-Flower and Ridley, 2015; in mammals, Rus Hoelzel et al., 1999; Ortega et al., 2003). In halfbeaks, dominant males have greater access to females and spend more time courting females than subordinate males. Since courtship effort is often positively correlated with male fitness (e.g., Shamble et al., 2009; Stoltz et al., 2009; Alonso et al., 2010), dominant male halfbeaks may increase their reproductive potential at the expense of subordinate males. The reproductive potential of socially subordinate male halfbeaks may be further impaired by the physiological costs associated with male-male competition. Specifically, Reuland et al. (2021) found that high levels of male-male competition led to impaired ejaculate quality in subordinate males. Such costs of social subordination are common in animals, with subordinate males often having higher levels of stress hormones (e.g., Fox et al., 1997; Creel, 2001; Milewski et al., 2022), lower levels of sex hormones (e.g., androgens, Oliveira et al., 2002; Nelson, 2005), and reduced investment in testicular tissue (e.g., Fitzpatrick et al., 2006). However, maintaining social dominance can be costly, potentially leading to trade-offs with other sexually selected or life history traits. (Fitzpatrick et al., 2012; Parker et al., 2013; Lüpold et al., 2014; Simmons and Emlen, 2006). Although our experimental time frame was too short to reveal the potential costs of social dominance, this remains an interesting avenue for future investigation. It is also interesting to note that even though the interaction between dominance and trial type was non-significant in our combined model, the separate models for each trial type indicate that the difference in courtship effort between dominant and subdominant males was lesser in trials without male-male competition (Table A1). This suggests that persistent physical interactions play a part in socially dominant male halfbeaks maintaining their advantage over subordinate males.

We found no evidence that male social dominance influences pre-copulatory female mate choice in halfbeaks, contradicting the general assumption that traits involved in male–male competition serve as honest signals for female choice (Berglund et al., 1996). Females did not alter their preference for dominant males after observing male–male interactions, and female association time with stimulus males was not predicted by male social dominance or by morphological differences between males. This finding contrasts with abundant empirical evidence showing that females often prefer dominant males as mates (Cox and Le Boeuf, 1977; Candolin, 1999; Kunc et al., 2006; Aquiloni et al., 2008; Aquiloni and Gherardi, 2010; Loranger and Bertram, 2016). Our results therefore suggest that female halfbeaks may be indifferent to male social dominance, even after having the opportunity to eavesdrop on male–male interactions. One possible explanation is that male-male competition only influences female preferences when it provides net fitness gains (Wong and Candolin, 2005). If dominance is not directly correlated to offspring viability, female halfbeaks may not gain useful information that will impact their fitness prospects from observing male bouts. For example, in the Pacific blue-eye fish *Pseudomugil signifier*, hatching success of eggs guarded by dominant or subordinate males did not differ, and females consequently showed no preference for winners or losers when privy to male competition (Wong, 2004). Similarly, in the sand goby *Pomatoschistus minutus*, females did not prefer dominant males but instead preferred good fathers (Forsgren, 1997). These results suggest that alternative traits may therefore be more important in signaling genetic quality or male investment in activities (e.g. defence of spawning sites) that translate into actual fitness gains to females. The potential links between social dominance and aspects of male quality that could offer females direct or indirect benefits in halfbeaks remain poorly understood and require further investigations.

An alternative explanation for our observed lack of female preference for dominant males could be that females are not under selection to discriminate based on male social dominance because male-male competition effectively eliminates subordinate males as potential suitors. This possibility was clearly evident from our experiment that allowed male-male competition during male-female interactions. By displacing subordinate males, dominant males may “pre-select” potential mates, resulting in females being predominantly courted by dominant males, and eliminating the need for females to exhibit an active preference for male social status (Wong and Candolin, 2005). Halfbeaks live in large mix sexed shoals where male-male agonistic behaviours are common (Devigili et al., 2021), suggesting that male-male social interactions mediate access to mates in wild halfbeak populations. This raises the possibility that strong male-male competition could relax selection on pre-copulatory female mate choice.

In conclusion, our results suggest that male–male competition, rather than pre-copulatory female preference, is the primary mechanism underlying the greater access to females exhibited by dominant male halfbeaks. Although females are often assumed to use the outcome of male-male competition to select high-quality mates (Berglund et al., 1996; Wiley and Poston, 1996; Wong and Candolin, 2005), our study challenges this assumption, since female halfbeaks showed no particular preference for males they observed winning intra-sexual contests. To prioritize tractability, our experimental design focused on the minimum number of males necessary to generate a dominance hierarchy: two males. However, in the wild halfbeaks live in shoals of 6 to 120 individuals (Devigili et al., 2021), where dominance effects may not scale linearly with group size. In larger groups, dominant males may face greater challenges in monopolizing mates (Newton-Fisher et al., 2010; Lardy et al., 2012), potentially weakening or altering the role of social dominance in securing male fitness (Reeve et al., 1998; Clutton-Brock, 1998). Future research should assess how male-male competition influences male fitness in more dynamic and ecologically relevant social conditions.

## Data availability

Data and scripts are available for reviewers as supplementary material.

## Appendix A

**Table A1:**
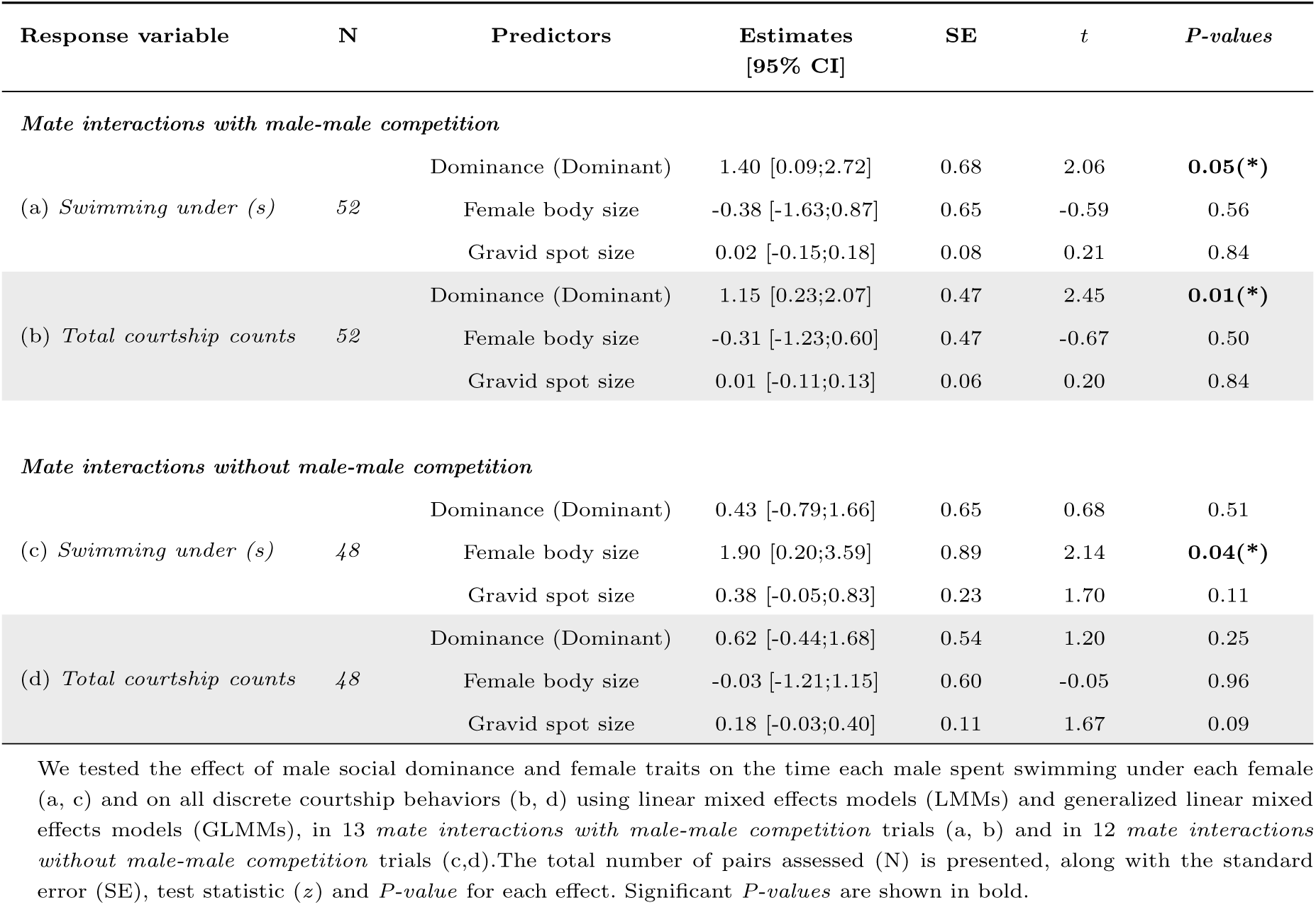
Effect of male social dominance and female traits on male access to females.

